# Validation of hypermethylated DNA regions found in colorectal cancers as potential aging-independent biomarkers of precancerous colorectal lesions

**DOI:** 10.1101/2023.05.24.542159

**Authors:** Sija Sajibu, Emanuel Sonder, Amit Tiwari, Stephany Orjuela, Hannah R. Parker, Olivier The Frans, Christoph Gubler, Giancarlo Marra, Mark D. Robinson

## Abstract

**Background:** We previously identified 16,772 colorectal cancer-associated hypermethylated DNA regions that were also detectable in precancerous colorectal lesions (preCRCs) and unrelated to normal mucosal aging. We have now conducted a study to validate 990 of these differentially methylated DNA regions (DMR) in a new series of preCRCs.

**Methods:** We used targeted bisulfite sequencing to validate these 990 potential biomarkers in 59 preCRC tissue samples (41 conventional adenomas, 18 sessile serrated lesions), each with a patient-matched normal mucosal sample. Based on differential DNA methylation tests, a panel of (candidate) DMRs was chosen on a subset of the (our) cohort and validated on the remaining part of our cohort and (two) further publicly available datasets with respect to their stratifying potential between preCRCs and normal mucosa.

**Results:** Strong statistical significance for the difference in methylation levels was observed across the full set of 990 investigated DMRs. From these, a selected candidate panel of 30 DMRs correctly identified 58/59 tumors (area under the receiver operating curve: 0.998).

**Conclusions:** These validated DNA hypermethylation markers can be exploited to develop more accurate noninvasive colorectal tumor screening assays.

## Background

The past 40 years have witnessed considerable reductions in both the mortality and incidence of colorectal cancer (CRC). These trends are largely attributable to the increasing use of population-based colonoscopy screening, which allows the identification and removal of early-stage CRCs as well as neoplastic lesions regarded as precancerous (preCRCs), such as conventional adenomas (cADNs) and sessile serrated lesions (SSLs) [1]. The presence of preCRCs precedes CRC onset by ∼10 years, thereby providing a substantial time window for preventive interventions [2].

Due to its high diagnostic efficacy, colonoscopy is now the primary tool for CRC screening, but it has several well-known shortcomings. Studies based on same-day tandem colonoscopy findings indicate that endoscopists overlook around 26% of adenomas [3], and missed precancerous lesions have been estimated to give rise to half of the CRCs detected within two years of a negative colonoscopy [4]. Furthermore, colonoscopy is relatively invasive and in rare cases associated with intestinal bleeding or perforation [2]. Finally, many patients consider the procedure and the bowel prep that precedes it uncomfortable, time-consuming, and/or generally “unpleasant.” Consequently, CRC screening strategies increasingly include a preliminary non-invasive testing step aimed at improving adherence to screening programs and obtaining additional information that can complement endoscopic findings.

Interest is growing in the development of DNA methylation-based markers for CRC screening. Compared with genetic mutations, methylation abnormalities are far more frequent in colorectal tumor genomes of all stages [5]. Although DNA methylation markers of CRC have been studied extensively, converting the findings into clinically useful tests has proven to be challenging [6,7]. The non-systematic selection of the candidate loci in these studies and the limited number of candidates subjected to testing are key factors in these difficulties [8].

Our group recently demonstrated that the abnormal methylation phenotype of specific DNA regions described in CRCs [5] is already evident during their precancerous stages [9]. We identified ∼ 20,000 CRC-associated hypermethylated DNA regions that also contained stretches of methylated CpG island-related cytosines in preCRCs—elements that were almost invariably unmethylated in tissue donor-matched samples of normal colon mucosa (NM) [9]. These findings suggest that a diagnostic test capable of identifying the presence of these differentially methylated regions (DMRs) (i.e., hypermethylated in tumors vs. NM) could improve the detection of both CRCs and their precancerous counterparts.

Some regions that were hypermethylated in preCRCs, however, also displayed mildly increased methylation in NM samples from older (vs. younger) tumor-free women. We therefore conducted a follow-up study to explore the overlap between the ∼20,000 tumor-associated DMRs and genomic regions that appeared to undergo methylation as an effect of aging (aging-associated DMRs) [10]. The vast majority of the tumor-associated DMRs (n=16,772) showed no methylation changes in the aging NM and were thus considered likely to be “*tumorigenesis-specific*” [10]. In the study described below, we selected a subset of these putatively high-potential age-independent markers of colorectal tumorigenesis for validation in a new series of prospectively collected colorectal tissues.

## Methods

### Tissue samples

DMR validation was performed on a series of colorectal tissue samples collected during colonoscopy. 59 preCRCs (41 cADNs, 18 SSLs), each with a patient-matched NM sample collected >2 cm from the tumor were included in the study (**Supplementary Table 1)**. Immediately after collection, each sample was immersed in AllProtect Tissue Reagent (Qiagen, Hilden, Germany), held overnight at 4°C, and stored at −80°C. DNA was extracted with the Qiagen AllPrep DNA/RNA mini kit (Hilden, Germany).

### Selection of DMRs to be validated in this study

**Supplementary Table 2** shows the 1096 genomic loci included in our validation set. These included 990 potential biomarkers of colorectal tumorigenesis drawn from the list of 16,772 DMRs we had previously classified [9,10] as “tumorigenesis-specific,” i.e., those displaying evident hypermethylation (vs. NM) in cADNs, SSLs, or both preCRC types but no methylation changes in NM that could be attributed to aging [10].

All 990 of the potential biomarker DMRs we selected had been identified in *both* cADNs and SSLs. The vast majority (935/990) also fulfilled all the following formal inclusion criteria: overlap with a CpG island; a length of >=80bp; a difference between preCRC and NM methylation levels of >=0.5 (methylation level range: from 0 to 1); and a q-value <0.05. The remaining 55 candidates were subjectively chosen on the basis of visual inspection of our raw data in the Integrative Genome View (IGV). While these loci were also markedly hypermethylated in both preCRC types, they failed to satisfy one or more of the formal inclusion criteria mentioned above.

For control purposes, the validation set also contained: 39 DMRs that had displayed substantial tumorigenesis-specific hypermethylation in only one type of preCRC (17 in cADNs, 22 in SSLs) [9, 10]; forty-seven genomic loci that had displayed high-level methylation in preCRCs *as well as* in donor-paired NM samples (referred to hereafter as “negative controls”); and 20 other DMRs we had classified as “aging-specific” (based on their absence in cADNs and SSLs and the presence in NM samples from tumor-free women aged 40-70 but not in NM samples from tumor-free women aged <40) (Supplementary Table 2).

### Probe design

Target enrichment of the selected genomic regions was achieved by solution-based hybrid capture performed with a synthetic library ultimately consisting of 96,456 RNA probes (myBaits®, Arbor Biosciences, Ann Arbor, MI, USA). Capture oligonucleotides (length: 80 nucleotides, tiling density: 2x) were designed to target approximately 0.5 Mb of the genome and contained 44,177 CpG sites derived from the Hg19 genome build. For each DMR, we designed nine probes for the detection the following methylation patterns: 1) genomic reference (genomic sequence before bisulfite conversion); 2) all CpG sites methylated in both cis and trans sequences; 3) no methylation in either cis or trans sequences, 4) a random half of the CpG sites methylated in both cis and trans sequences, and 5) the other half of the CpG sites methylated for both cis and trans sequences. (See Supplementary Table 2 for the genomic coordinates of these probes.)

### Library preparation and target enrichment

Genomic DNA was extracted from colorectal tissues using the AllPrep DNA/RNA kit (Qiagen, Hilden, Germany). DNA concentrations were quantified with a Qubit fluorometer (Invitrogen, Carlsbad, CA, USA), and quality was assessed with a NanoDrop spectrophotometer (Thermo Fisher Scientific, Wilmington, NC, USA). Genomic DNA (100ng) was bisulfite-converted with the EZ-DNA Methylation-Gold kit (Zymo, Irvine, CA, USA) prior to preparation of the library with the Accel-NGS Methyl-Seq DNA Library Kit (Swift Biosciences, Ann Arbor, MI, USA). Equimolar amounts of eight indexed libraries were pooled for hybridization with customized RNA baits, which was performed twice at 63°C for 16 to 24 hours. AMPure Xp beads (Beckman Coulter Inc, Brea, CA, USA) were used to isolate biotinylated DNA, and the bead-bound enriched libraries were amplified with KAPA HiFi HotStart Ready Mix (KAPA Biosystems, Wilmington, MA, USA) and adaptor-specific primers. The concentrations of the enriched libraries were measured with a Qubit fluorometer. Library size was determined with a Tapestation (Agilent, Santa Clara, CA, USA) for subsequent library titration and next generation sequencing. Each pooled library was sequenced on an Illumina NovaSeq 6000 instrument (San Diego, CA, USA) using one lane to generate an average of 25 million (2 x 150bp) reads per sample.

### Data analysis

The patient sample database was split by batch (i.e., date of measurement) to obtain a *training set* consisting of 70 samples (35 patients, 7 batches) and a *test set* comprising 48 samples (24 patients, 3 batches). Per-CpG differential methylation (DM) tests were conducted in a per-patient paired design using the limma R package. Per-region p-values were obtained by aggregating all per-CpG p-values using the Simes Method [11]. A candidate biomarker panel was then created that included the 30 DMRs with the lowest p-values in the training set. This panel (referred to hereafter as the *top 30 candidate biomarker panel*) was then further validated in the test set.

### Preprocessing

Paired-end sequencing reads were trimmed using Trim Galore (0.6.5) software [12] and subsequently aligned using Bismark (0.23.0) [13] with Bowtie2 (2.3.5.1) [14] to the GRCh37/hg19 human reference genome. Deduplication was performed with the Bismark deduplication functionality, and cytosine methylation calls were obtained with the Bismark Methylation Extractor. Deduplicated per-CpG reports were imported and handled using the R bsseq library [15] and the hg19 BS-genome reference. Of the 44,177 CpGs that were probed, 43,786 were covered (by at least seven reads) in all the samples and therefore considered for subsequent analysis.

### Training/Test split

For the selection of a top 30 DMRs, we split the patient population (n=59) by batch (i.e., the date of measurement) in a stratified manner, considering the ratio of SSL and cADN samples per-batch using the function stratified from the splitstackshape package [16].

### Selection of DMRs for the top 30 candidate biomarker panel

The training dataset was subjected to differential methylation (DM) testing to select the regions characterized by consistent methylation differences between precancerous colorectal lesions (preCRCs) and normal colon mucosa (NM). The DM-strategy chosen was based on functionalities of the limma R package [17,18, 19], which allow for robustified estimates of the degrees of freedom (df) and variance priors in the presence of outliers.

Moderated t-statistics based on log2CPM-transformed read counts were obtained per-CpG using the modelMatrixMeth function from the edgeR package [20] for computing the design matrix, accounting for the two-level factor read count structure of DNA methylation data, thereby loosely following the rationale of the per-CpG test implemented in the DMRcate package [21]. All tests were conducted in a per-patient paired design. Per-CpG p-values were aggregated for each region using the Simes method [11]. P-values were corrected for multiple testing using the Benjamini and Hochberg method [22] in the stats R package [23]. For the panel, the 30 regions with the lowest corrected p-values were selected.

### Validation

To test the stratifying potential of the top 30 candidate biomarker DMRs, we calculated the mean methylation level of the regions across the panel for each sample. In addition to the test set described above, we used two published datasets, each comprising a sufficiently high number of colorectal tissue samples and NM samples. The first (referred to hereafter as the GSE131013 dataset) [24] was generated in a 450K microarray-based study and includes methylation data on 90 donor-matched pairs of preCRC and NM tissue from patients and 48 samples of NM from healthy donors. The second (referred to hereafter as the GSE48684 dataset) [25] was generated with the Illumina methylation 450K microarray and comprises methylation data on 42 cADNs, 64 CRCs, and 41 unmatched NM tissues.

For the GSE131013 and GSE48684 datasets from the literature, we considered the Beta-values of all CpGs overlapping the region coordinates of this study (as downloaded May 2022) for the subsequent tasks. Receiver operating characteristics (ROC) curves were obtained, and the area under the curve (AUC) for each dataset was calculated using the auc function from the pROC package [26]. Tissue samples were binarily labeled “NM” or “non-NM” (the latter comprising all preCRC and CRC tissues).

The 100 randomly selected sets of 30 DMRs each shown in the ROC plot were taken from the 990 candidate biomarker DMRs, and the mean methylation level was used to calculate the ROC curves.

The F1-score was applied using the cutpointr [27] R package to find a cutoff for the average per-region methylation level of the candidate marker panel that differentiates between preCRC and NM tissues in the training samples. Methylation levels equal to or exceeding the optimal cutoff were considered to originate from preCRC lesions. Originally, a procedure giving a 2-, 5-, or 10-times higher weight to false negatives than to false positives was considered on the training dataset. However, given the limited size of that dataset, this would have resulted in cutoffs taking infinite values and therefore in all samples being regarded as positives. Hence, the F1-score was used on the training dataset to identify a cutoff. In fact, the optimal cutoff identified in this manner was concordant with that identified using Youden’s J statistic or Cohen’s Kappa.

### Multidimensional scaling (MDS) plots

All MDS plots were generated using asin-transformed per-region methylation levels. Subsets of the regions (i.e., the top 100 variable regions for each pair of samples) were used to determine the MDS dimensions with the plotMDS function [17].

### Heatmaps

The heatmaps showing methylation differences reflect the difference between the methylation level per region (i.e., the fraction of methylated over all reads) as obtained by the getMeth(…,what=perRegion”) function of the bsseq library.

## Results

The approach used to validate the 990 DMRs in our new series of preCRC tumor tissues was similar to that used in our earlier studies [9,10]. Because the Roche SeqCap Epi CpGiant kit used in those studies had been discontinued, we used a custom-designed set of probes. This restricted our methylation analysis to approximately 1000 genomic regions (as compared with the ∼16,000 analyzed with the Roche kit (see *Methods*) and reduced the amount of DNA needed per sample from 1000ng to 100ng—a substantial advantage for future studies, especially those conducted on low-volume tissue samples. We comparatively analyzed two preCRC lesions and their matched NM samples with the custom *myBaits* protocol and the previously used SeqCap Epi CpGiant protocol. The strong correlation (r=0.97) observed between methylation values obtained with the two methods (**Figure 1**) indicate that the protocols are equally robust.

**Figure 1:**
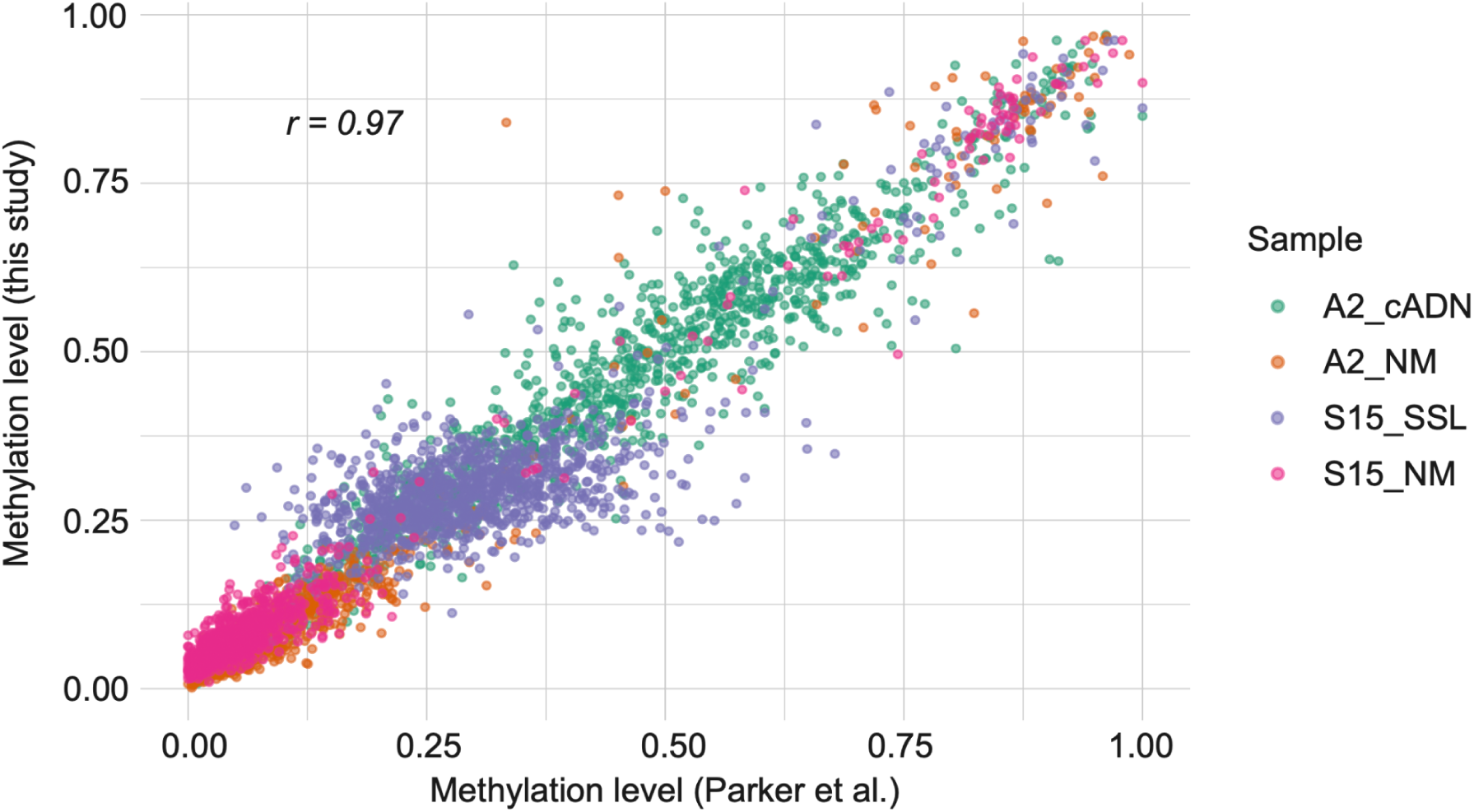
Comparison of the methylation data obtained with the protocol used in this study and in our previous study. [9]. Correlation plot (Pearson correlation r) of the per-region averaged methylation levels for the four samples (color-coded by tissue of origin) profiled with both protocols. Each dot represents a genomic locus validated in this study.

The DNA methylomes of cADNs and SSLs were clearly distinguished from those of their matched NM samples (**Figures 2A** and **B**). As expected, the DMRs classified in our earlier studies as cADN-specific or SSL-specific displayed similar tumor-specificities in this validation study, and the methylation statuses of the negative-control and aging-specific DMRs were consistent across all samples (**Figure 2B**). All 990 candidate biomarker DMRs performed well in the training dataset (24 cADNs and 11 SSLs), with very low false discovery rates (FDRs) (2.77E-20 to 5.24E-06) and substantial methylation changes (per-DMR mean log fold changes of CpGs 5.43 to 1.13, vs. NM) (**Supplementary Table 2**).

**Figure 2:**
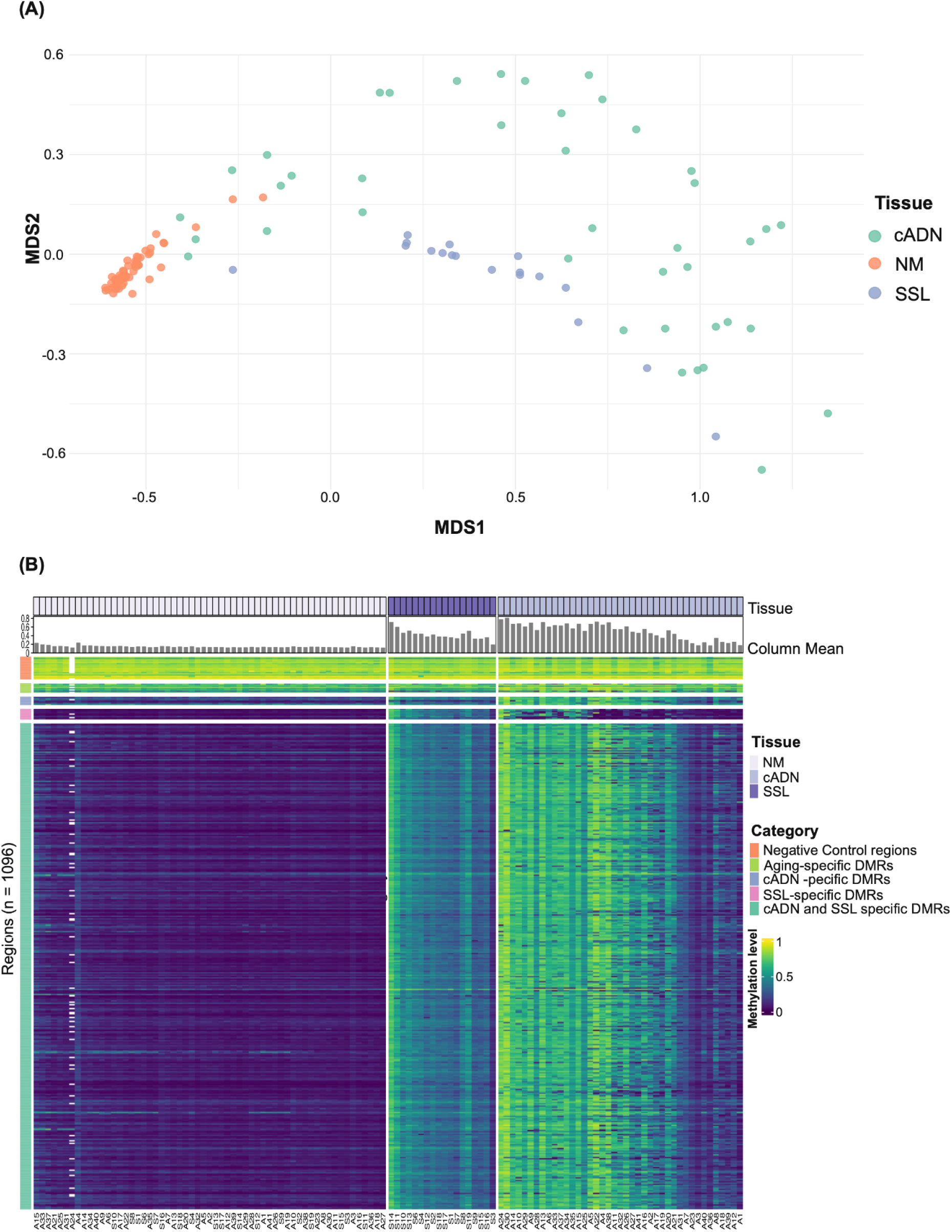
Unsupervised analysis of the methylation levels in the 118 tissue samples investigated. **(A)** Multidimensional scaling (MDS) plot of the samples (cADNs n=41, SSLs n=18 and matched NM (n=59) based on DNA methylation levels in the 100 most variable DMRs. Per-DMR (averaged) methylation levels were subjected to asin/arcsin transformation. **(B)** Heatmap visualization of mean methylation levels averaged across all CpG sites of each of the 1096 genomic regions analyzed in each tissue sample (numerically coded and prefixed with “A” for cADN or “S” for SSL). In both lesion types, methylation levels at the 990 candidate biomarker DMRs were generally higher than those in their matched NM samples. In contrast, the preCRC and NM methylation profiles for the negative-control and aging-specific DMRs were similar. Column Mean: Mean methylation level per sample (averaged across CpG sites of the 990 candidate biomarker DMRs). White blocks: genomic regions not covered by any reads in the A24 cADN tissue sample.

We then validated the top 30 candidate biomarker DMRs using our own test dataset and two published GSE131013 [24] and GSE48684 [25] datasets (*Methods*): As shown in Figures 3A and **B**, all the tumors in our training and test sets clearly displayed some degree of hypermethylation at all 30 of these loci. In these two datasets, the Top 30 Candidate marker panel with a mean methylation level cut-off of 0.0918 (methylation level range: 0 to 1) correctly identified 58 (98.3%) of the 59 preCRCs as tumors (vs. NM), with one preCRC misclassified as NM in the test dataset. As shown in Figure 3C, the top 30 candidate marker panel distinguished almost all the tumors from NM samples with high accuracy (AUCs: 0.998, 0.981, 0.977 in our test dataset, GSE131013, and GSE48684 respectively). Panels composed of 30 cADN-& SSL-specific DMRs randomly selected from the 990 shown in Supplementary Table 2 (gray curves in Figure 3C) also performed very well across our test dataset (AUC: 0.994) and satisfactorily across the external datasets (AUC: 0.971 and 0.962 in GSE131013 and GSE48684, respectively). Across the three datasets, the average performance of these randomly selected panels was superior to those of methylation markers used in the currently available CRC screening tests (e.g., *VIM*, *NDRG4/BMP3*, and *SEPT9*; see *Discussion*).

**Figure 3:**
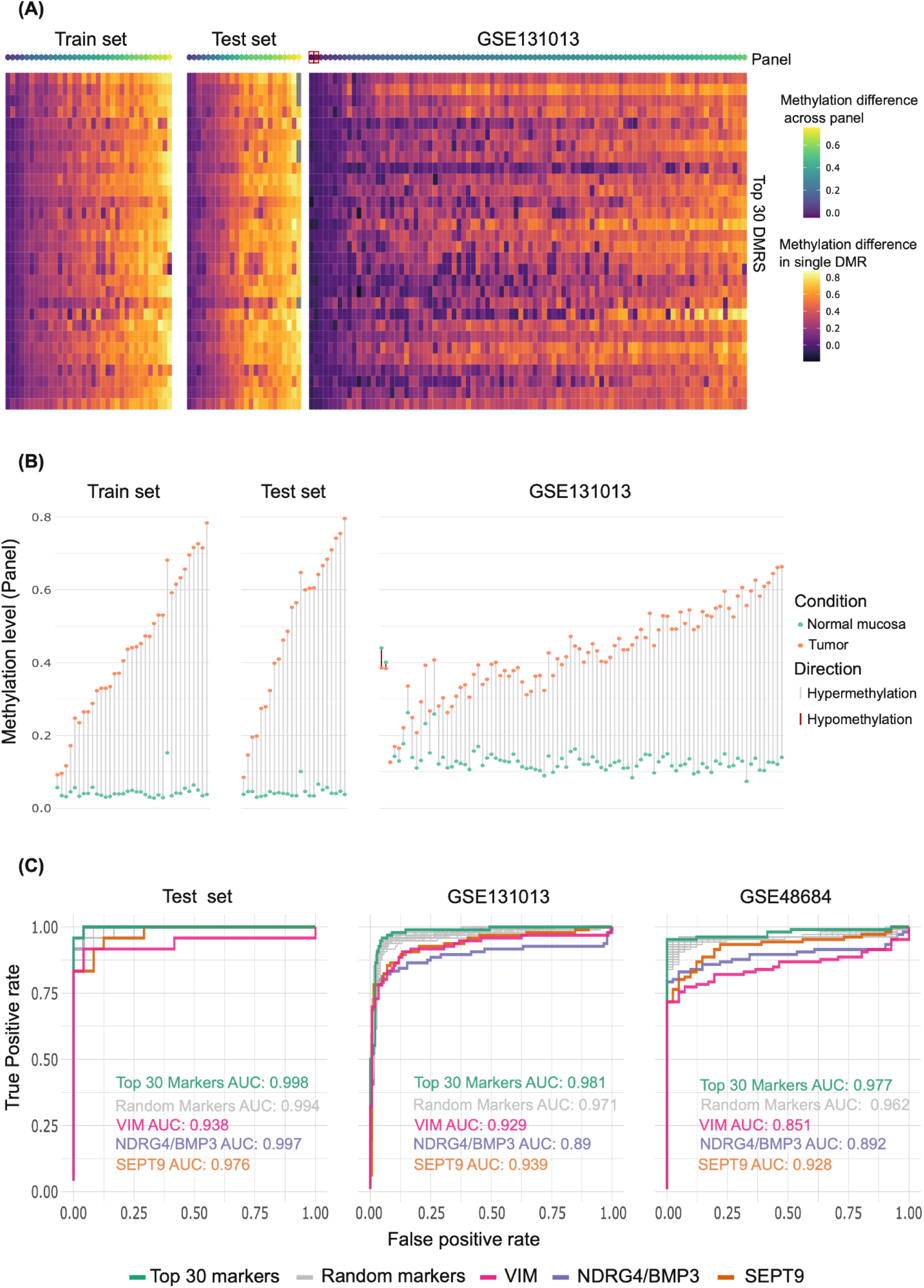
Validation of the top 30 candidate markers from Supplementary Table 2. (**A**) Heatmap showing the difference between per-DMR averaged methylation levels in normal mucosa (NM) and tumor tissues in our Training Set (70 samples from 35 patients), our Test Set (48 samples from 24 patients), and the GSE131013 tissue set (180 samples from 90 patients) (see *Methods*). The top row (“Panel”) shows the average methylation difference per patient across the 30 DMR marker candidates. (Red squares in this row: donors with hypomethylated tumors reflected by a negative average difference between their tumor and NM methylation levels.) Pairwise comparison of methylation levels in the GSE48684 samples could not be performed because per patient paired sample data was not available (see *Methods)*. (**B**) Averaged methylation levels for the top 30 marker candidates in tumors (orange dots) and paired NM (green dots) for each patient in the three datasets mentioned in (A). (Two tumors of the GSE131013 dataset displayed hypo-rather than hypermethylation relative to their matched NM samples.) (**C**) Receiver operating characteristic (ROC) curves showing the differentiating potential of the “per sample mean methylation level” at our top 30 candidate marker DMR set, at the 100 randomly selected sets of 30 DMRs from our Supplementary Table 2, at *VIM* DMR, at *NDRG4/BMP3* DMR and at *SEPT9* DMR in our test dataset and the GSE131013 and GSE48684 datasets. AUCs are given in each graph (the AUC of the randomly selected sets of 30 DMRs, i.e., gray curve, represents the mean of AUCs obtained across all 100 random sets).

## Discussion

With this validation study conducted on a new prospectively collected series of colorectal tissues, we have demonstrated that numerous hypermethylated DMRs are robust indicators of the precancerous stages of colorectal tumorigenesis. Because these same DNA regions are also highly hypermethylated in CRC [9, 10], they have the potential for development as screening biomarkers for colorectal tumors of all stages. Several studies have already reported DNA methylation alterations in CRC, proposing their use as a diagnostic marker [5]. Few studies have been performed in precancerous lesions, generally in cADNs and few or no SSLs [24, 25, 28]. In our previous studies [9, 10] and in the present study, we have analyzed DMRs developing in both cADNs and SSLs (i.e., the vast majority of precancerous colorectal lesions). Therefore, these epigenetic markers can effectively detect colorectal tumors developing along either the adenomatous or the serrated pathway of tumorigenesis, and they tend to be found across most lesions unlike mutations (e.g., those involving *KRAS* or *BRAF*, which are identified in only ∼40% and ∼10% of colorectal neoplasms, respectively).

Pre-colonoscopic screening assays based on the methylation analysis of fecal DNA might thus complement (or even replace) the immunochemical stool tests widely used in today’s clinics for the identification of individuals who might need colonoscopy. Indeed, some currently available fecal DNA assays include one or two methylation markers, such as hypermethylation of *VIM* (ColoSure assay) or *NDRG4* and *BMP3* (Cologuard assay) [29], and the blood DNA-based assay Epi proColon tests for hypermethylation of *SEPT9* [30]. At present, none of these assays are widely used in clinical practice, mainly because they have not been proven to be superior to currently-used immunochemical tests in detecting CRC. As for their accuracy in detecting preCRCs, it is far from optimal (11.2% to 42.4% sensitivity; 86.6% to 91.5% specificity) [29,30]. Our top 30 candidate DMRs performed better, in our test set and in external datasets, than the DMRs included in these commercial assays, suggesting that a larger set of reliable DNA methylation markers (e.g., 30 cADN-and SSL-specific, aging-independent DMRs) might allow more sensitive and more specific detection of precancerous lesions. Therefore, the set of DMRs we have investigated seems highly suitable for developing novel, noninvasive screening tests for the early detection of all colorectal tumors. Prospective studies in stool-DNA samples should be carried out to verify the clinical potential of our markers.

Our findings also indicate that a screening assay based exclusively on DNA hypermethylation markers is likely to miss a few preCRCs (∼2% in our validation study, see *Results*) owing to their very low levels of DNA hypermethylation. A more promising strategy for enhancing assay sensitivity might therefore involve adding methylation-based markers to panels containing immunochemical and genetic markers, an approach similar to the one used in the Cologuard assay.

## Conclusions

We have validated an extensive set of DMRs as markers capable of accurately predicting the presence of a colorectal tumor regardless of the patient’s age. The list of DMRs validated in this study can thus be productively mined to establish promising biomarker sets for fecal DNA based screening assays.

### List of abbreviations

AUC: area under the curves
cADN: conventional adenomas
CRC: colorectal cancer
DF: degrees of freedom
DM: differential methylation
DMR: differentially methylated region
FDRs: false discovery rates
IGV: Integrative Genome View
MDS: Multidimensional scaling
NM: normal colon mucosa
preCRC: precancerous colorectal lesion
ROC: receiver operating characteristic
SSL: sessile serrated lesion

## Declarations

### Ethical approval and consent to participate

The study was approved by Zurich’s Cantonal Ethics Committee, Switzerland (KEK-ZH-Nr: 2015-0068 and BASEC-Nr.: 2015-00185). All methods were carried out in accordance with relevant guidelines and regulations. Donors provided written informed consent to tissue collection, testing, and data publication. Samples were numerically coded to protect donor rights to confidentiality and privacy.

## Consent for publication

Not Applicable.

## Availability of data and materials

The dataset supporting the conclusions of this article is available in the ArrayExpress repository under the accession number E-MTAB-12612. Supplementary table 2 has been deposited into the Zenodo open data repository: DOI: 10.5281/zenodo.7715366 (https://doi.org/10.5281/zenodo.7715366). All code used for the analysis can be found under the following repository link: https://github.com/emsonder/validation_dmrs (DOI: 10.5281/zenodo.7704489, https://doi.org/10.5281/zenodo.7704489)

## Authors’ contributions

G.M., M.D.R., and S.S. designed the research; S.S. and H.R.P. processed tissue samples and conducted experiments for DNA methylation analysis; O.T.F. and C.G. performed tissue sample and clinical data collection; E.S., S.O. and A.T. analyzed data; S.S., E.S., A.T., and G.M. wrote the paper.

## Acknowledgments

We thank Catharine Aquino, Andrea Patrignani, and Luca Albanese for technical support; Federico Buffoli, Fabrizio Cereatti, and Matthias Sauter for help in previous studies that made the present one possible; and Marian Everett Kent for editing the manuscript.

## Supplementary Tables

**Supplementary Table 1**: Characteristics of the 59 precancerous lesions used in the study.

**Supplementary table 2**: List of the 1096 genomic loci investigated in the study (ordered by lowest corrected q-values). Genome coordinates are based on the GRCh37/hg19 genome assembly.

